# DeepPWM-BindingNet: Unleashing Binding Prediction with Combined Sequence and PWM Features

**DOI:** 10.1101/2024.08.21.609040

**Authors:** Sarwan Ali, Prakash Chourasia, Murray Patterson

**Author notes:** {, }. Equal Contribution (Sarwan Ali, Prakash Chourasia).

## Abstract

A crucial challenge in molecular biology is the prediction of DNA-protein binding interactions, which has applications in the study of gene regulation and genome functionality. In this paper, we present a novel deep-learning framework to predict DNA-protein binding interactions with increased precision and interoperability. Our proposed framework DeepPWM-BindingNet leverages the rich information encoded in Position Weight Matrices (PWMs), which capture the sequence-specific binding preferences of proteins. These PWM-derived features are seamlessly integrated into a hybrid model of convolutional recurrent neural networks (CRNNs) that extracts hierarchical features from DNA sequences and protein structures. The sequential dependencies within the sequences are captured by recurrent layers. By incorporating PWM-derived features, the model’s interpretability is improved, enabling researchers to learn more about the underlying binding mechanisms. The model’s capacity to locate crucial binding sites is improved by the incorporation of an attention mechanism that highlights crucial regions. Experiments on diverse DNA-protein interaction datasets demonstrate the proposed approach improves the predictive performance. The proposed model holds significant potential in deciphering intricate DNA-protein interactions, ultimately advancing our comprehension of gene regulation mechanisms.

## 1 Introduction

The interaction between DNA and proteins is pivotal in various cellular processes, including gene expression, DNA repair, and signal transduction. Accurate prediction of DNA-protein binding sites is essential for understanding the molecular mechanisms that govern these processes. Other applications include drug discovery [1], understanding gene regulation [2], and disease prediction [3]. Deep learning techniques have shown promise for sequence analysis, as they can automatically learn complex patterns from DNA and protein sequences [4,5,6,7]. With the advent of high-throughput sequencing technologies, there is an increasing demand for computational models that can effectively predict binding interactions between DNA sequences and proteins [8].

Traditional approaches to DNA-protein binding prediction have relied on sequence analysis tools like MEME (Multiple Em for Motif Elicitation) [9] and Gibbs Motif Sampler [10] are used to identify DNA motifs that are likely binding sites for specific proteins. Tools like TRANSFAC [11] and JASPAR [12] provide databases of known transcription factor binding motifs. Also, statistical methods like position weight matrices (PWMs) can be employed to predict binding sites based on the frequency of nucleotides at each position in a set of known binding sites [13]. ChIP is a technique used to identify DNA sequences associated with specific proteins, often transcription factors [14]. It involves cross-linking DNA and proteins, immunoprecipitating the protein of interest, and then sequencing the associated DNA fragments to determine binding sites [15]. However, these methods often struggle to capture the intricate sequence patterns contributing to binding specificity [16]. Moreover, these methods can be costly due to the need for specialized equipment, reagents, and expertise [16]. While bioinformatics tools can assist in analyzing experimental data, they often require prior knowledge of binding motifs or binding partners [3]. Identifying novel motifs or interactions may be difficult with these methods. Some methods, like EMSA [17], ChIP [14], and footprinting, [18] may lack the sensitivity to detect weak or transient interactions [8]. Additionally, they may not provide high specificity, leading to false positives or negatives. These methods are typically limited to studying interactions with known proteins or transcription factors. Identifying novel binding partners can be challenging using these approaches.

Deep learning has emerged as a powerful tool for capturing complex relationships within biological data in recent years. Convolutional neural networks (CNNs) and recurrent neural networks (RNNs) have demonstrated impressive performance in various bioinformatics tasks, including DNA-protein binding prediction [4]. More complex techniques may anticipate binding locations with greater accuracy [19]. Deep learning has recently been effectively used in several fields and demonstrated some excellent performance, including motif discovery [4]. Modeling the sequence peculiarities of DNA-binding proteins using deep convolutional neural network (CNN), a variant of multilayer artificial neural network specialized for processing images, was accomplished by DeepBind [20], and its performance is superior to some best existing conventional methods. Existing deep learning techniques have excelled lately, but they still have certain drawbacks. They overlooked the high-order correlations among nucleotides in practice and only took into account the independent interaction among nucleotides in the binding sites [21]. Also, they merely used a set motif length to identify the binding characteristics in the genomic sequences.

We introduce DeepPWM-BindingNet, a novel deep-learning architecture tailored for DNA-protein binding prediction. Our approach leverages both the sequence information encoded in DNA sequences and protein structures and the domain-specific knowledge captured by Position Weight Matrices (PWMs). PWMs are widely used in bioinformatics to represent the binding preferences of proteins at different positions. PWMs represent the binding preferences of proteins at various positions, reflecting empirical data on these interactions. By integrating PWM-derived features with deep learning, we aim to enhance the accuracy and interpretability of DNA-protein binding predictions. The key contributions of this study are as follows:

1. Integration of PWM-Derived Features: We propose a novel approach that integrates PWM-derived features into a deep learning framework. These features provide crucial insights into the specific binding preferences of proteins at different positions along DNA sequences.
2. Hierarchical Feature Extraction: DeepPWM-BindingNet combines CNNs and RNNs to extract hierarchical features from DNA sequences and protein structures. This enables the model to capture local and global sequence patterns, contributing to improved predictive performance.
3. Attention Mechanism: To focus on critical regions within sequences, we incorporate an attention mechanism that allows the model to weigh the importance of different segments. This enhances the model’s ability to identify essential binding sites and improves interpretability.
4. Enhanced Predictive Performance: Through extensive experimentation on diverse DNA-protein interaction datasets, we demonstrate that DeepPWM-BindingNet outperforms existing methods in terms of predictive accuracy. The integration of PWM-derived features further boosts the model’s performance.
5. Interpretability: Integrating PWM-derived features provides researchers with a more interpretable model. This enables a deeper understanding of the underlying mechanisms driving DNA-protein binding interactions.

In the remaining sections, we provide a brief literature review in Section 2. Architecture and methodology of DeepPWM-BindingNet in Section 3. The details regarding the dataset statistics and baseline models are presented in Section 4. We then present experimental results and comparative analyses with existing methods in Section 5. Finally, the conclusion of the whole study is presented in Section 6.

## 2 Related Work

The prediction of DNA-protein binding interactions has been a subject of extensive research, with a wide range of computational methods developed over the years. These methods can be broadly categorized into sequence-based, structure-based, and hybrid approaches that integrate sequence and structure information.

Early approaches relied on identifying sequence motifs – short, conserved sequences – as indicative of binding sites. MEME (Multiple Em for Motif Elicitation) [9] and FIMO (Find Individual Motif Occurrences) [22] are widely used for motif discovery and binding site prediction. However, these methods often fail to capture subtle dependencies between nucleotides and amino acids, contributing to binding specificity. Structure-based approaches leverage protein structures to predict binding interactions [23]. Molecular docking methods [24], such as AutoDock and Rosetta [25], simulate the interaction between DNA and proteins to predict their binding affinity [26]. These models are computationally expensive and rely on accurate structural information [27]. Several recent authors proposed structure-based and sequence-based methods for sequence analyses [28,29,30].

Recent advancements in deep learning have spurred the development of hybrid methods that combine sequence and structure information for accurate binding prediction [31]. Methods like DeepBind [32] and DeepSEA [33] employ convolutional neural networks (CNNs) to learn sequence motifs and predict binding sites. LLM-based methods are used in [34]. While these methods have shown promising results, they often overlook the finer details of sequence-structure relationships. Integration of Position Weight Matrices (PWMs): In this study, we propose the integration of Position Weight Matrices (PWMs) – a well-established tool in bioinformatics – into a deep learning framework [35]. PWMs [36] represent the binding preferences of proteins at different positions along DNA sequences, capturing the specific nucleotide and amino acid interactions. We aim to bridge the gap between sequence and structure information by incorporating PWM-derived features, enhancing predictive accuracy.

## 3 Proposed Approach

In this section, we describe the proposed approach for predicting DNA-protein binding, called DeepPWM-BindingNet. DeepPWM-BindingNet capitalizes on the strengths of both deep learning and position weight matrix (PWM)-derived features. The architecture combines convolutional and recurrent layers to extract hierarchical features from DNA sequences and protein structures. This approach allows the model to learn local sequence patterns and global interactions, enhancing its predictive capabilities. Integrating an attention mechanism further refines the model’s predictions by focusing on crucial regions within sequences. This enables DeepPWM-BindingNet to identify essential binding sites and improve interpretability, a critical factor in biological research. The key steps of our approach are as follows:

### 3.1 Data Preprocessing

The first step in our approach involves data preprocessing to prepare the input data for the deep learning model.

– **Sequence Data**: We obtain DNA and protein sequence data, where each sequence is represented as a string of amino acids or nucleotides.
– **One-Hot Encoding**: We convert the sequence data into one-hot encoding. Each amino acid or nucleotide is represented as a binary vector, where each position in the vector corresponds to a specific amino acid or nucleotide. This encoding allows the model to process sequence data as numerical input.
– **Sequence Padding**: To ensure uniform input dimensions, we pad the sequences to a fixed length, typically achieved by adding zeros to sequences that are shorter than the maximum sequence length.
– **PWM Feature Extraction**: We compute Position Weight Matrices (PWM) for each sequence. PWMs capture positional information about nucleotide or amino acid frequencies in the sequences.
– **Normalization**: We normalize the PWM-derived feature vectors to have zero mean and unit variance to enhance model training.
– **Concatenation**: We concatenate the one-hot encoded sequences and the normalized PWM-derived feature vectors to create the final input dataset.

### 3.2 Model Architecture

Our deep learning model architecture is designed to learn features from the concatenated input data effectively.

– **Convolutional Layers**: We use one-dimensional convolutional layers to capture local patterns and features in the sequences. These layers consist of multiple filters with varying kernel sizes to capture different scale features.
– **Max-Pooling Layers**: Max-pooling layers follow the convolutional layers to downsample the feature maps, retaining the most relevant information.
– **Bidirectional LSTM Layer**: We employ a bidirectional Long Short-Term Memory (LSTM) layer to capture sequential dependencies and long-range interactions in the data. The bidirectional nature allows the model to consider both past and future contexts.
– **Attention Mechanism**: An attention mechanism is applied to the LSTM output, enabling the model to focus on specific parts of the sequence that are most informative for the prediction.
– **Global Average Pooling**: Global average pooling is performed on the attention-weighted LSTM output to reduce the spatial dimensions while retaining important features.
– **Dense Layers with Regularization**: We add densely connected layers with ReLU activation functions and L2 regularization to extract high-level features from the global average pooled output.
– **Output Layer**: The final output layer uses a softmax activation function for classification into binding/non-binding classes.

The detail regarding the architecture and number of parameters is reported in Table 1 for the PDB186 dataset (see Section 4.1 for details related to the dataset). In the model we have used *relu* activation function for regularization and used *softmax* for the output layer.

**Table 1:** Architecture and number of parameters for the proposed DeepPWMBindingNet model on PDB186 dataset (see Section 4.1 for details related to the dataset).

| Layer (type) | Output Shape | Param # | Connected to |
| --- | --- | --- | --- |
| input (InputLayer) | [(None, 1323, 42)] | 0 | - |
| conv1d (Conv1D) | (None, 1323, 64) | 18880 | input[0][0] |
| max_pooling1d (MaxPooling1D) | (None, 661, 64) | 0 | conv1d[0][0] |
| conv1d_1 (Conv1D) | (None, 661, 128) | 41088 | max_pooling1d[0][0] |
| max_pooling1d_1 (MaxPooling1D) | (None, 330, 128) | 0 | conv1d_1[0][0] |
| bidirectional (Bidirectional) | (None, 330, 128) | 98816 | max_pooling1d_1[0][0] |
| attention (Attention) | (None, 330, 128) | 0 | bidirectional[0][0] |
| - | - | - | bidirectional[0][0] |
| global_average_pooling1d (GlobalAveragePooling1D) | (None, 128) | 0 | attention[0][0] |
| dense (Dense) | (None, 128) | 16512 | global_average_pooling1d[0][0] |
| dropout (Dropout) | (None, 128) | 0 | dense[0][0] |
| dense_1 (Dense) | (None, 64) | 8256 | dropout[0][0] |
| dropout_1 (Dropout) | (None, 64) | 0 | dense_1[0][0] |
| dense_2 (Dense) | (None, 2) | 130 | dropout_1[0][0] |
| Total params: 183,682 | - | - | - |
| Trainable params: 183,682 | - | - | - |
| Non-trainable params: 0 | - | - | - |

### 3.3 Model Training

We train the deep learning model using the prepared dataset with the following configurations:

– **Loss Function**: Binary cross-entropy loss [37] is used for the classification (see Equation 1).
– **Optimizer**: We use the Adam optimizer to update model weights during training.
– **Callbacks**: Callbacks such as learning rate reduction and early stopping are employed to optimize training and prevent overfitting.
– **Batch Size and Epochs**: Training is performed in mini-batches with a specified batch size, and the process is repeated for a predefined number of epochs.

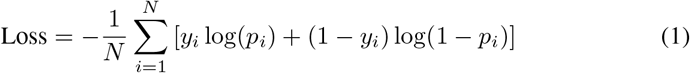

where y is the class label and *p* is the probability for prediction.

We assess the model’s performance using various evaluation metrics, including accuracy, precision, negative predictive value (NPV), sensitivity, specificity, Matthews correlation coefficient (MCC), F1-score, area under the ROC curve (AUC-ROC), and area under the precision-recall curve (AUC-PR).

To ensure robustness and reliability, we employ k-fold cross-validation, where the dataset is divided into k subsets (folds), and the model is trained and evaluated k times, with each fold serving as the test set once. In our experiments, we use *k* = 5 and report average ± standard deviation results for 5 folds.

## 4 Experimental Setup

In this section, we delve into the specifics of the dataset utilized in our experiments along with the details regarding the baseline methods used for results comparisons with the proposed approach. The experiments are conducted on a 64-bit Ubuntu operating system (version 16.04.7 LTS Xenial Xerus), utilizing a system equipped with an Intel(R) Xeon(R) CPU E7-4850 v4 clocked at 2.10GHz. The system boasted a substantial memory capacity of 3023 GB.

### 4.1 Dataset Statistics

We used the following benchmark datasets to perform experimentation. The statistics for all datasets discussed above are given in Table 2.

**Table 2:** Statistics for DNA-binding and non-binding protein sequences datasets.

| Datasets | Total Samples |  | Length Statistics |  |  |
| --- | --- | --- | --- | --- | --- |
|  | Negative | Positive | Min | Max | Mean |
| PDB14189 | 7060 | 7129 | 51 | 4911 | 425.313 |
| PDB2272 | 1119 | 1153 | 51 | 5183 | 459.907 |
| PDB1075 | 550 | 525 | 51 | 1323 | 240.213 |
| PDB186 | 93 | 93 | 64 | 1323 | 264.693 |

1. The PDB14189 dataset, obtained from [38], contains 14189 DNA-binding protein sequences (DBPs) and non-binding protein sequences (NDBPs). These sequences were collected from the UniProt database ^1^.
2. The PDB2272 dataset comprised of 2272 DBPs and NDBPs [39], originally obtained from Swiss-Prot ^2^. In this dataset, all proteins share sequence similarity of less than 25% with each other. As a preprocessing step, the sequences with irregular characters, e.g. “X” or “Z”, are removed.
3. The PDB1075 dataset, comprised of 1075 DBPs and NDBPs, is obtained from [40]. The number of positive samples (i.e. DBPs) in this dataset is 550 while the number of negative samples (i.e. NDBPs) is 525.
4. The final dataset that we used, called PDB186, comprised of 186 DBPs and NDBPs [41]. The number of positive samples (i.e. DBPs) in this dataset is 93 while the number of negative samples (i.e. NDBPs) is 93.

We have a balanced class for all our datasets as mentioned in table 2, which depicts that there is no bias for a certain value. As preprocessing we filtered out sequences with length where |*sequence*| < 50 and |*sequence*| > 6000.

### 4.2 Baseline Methods

To assess our proposed method, we compare it with the following methods:

#### MLapSVM [42]

The authors in [42] address the challenge of identifying DNA-binding proteins (DBPs), which are crucial in various cellular processes. They propose a novel method, called Multiple Laplacian Regularized Support Vector Machine with Local Behavior Similarity (MLapSVM-LBS), which combines three features extracted from protein sequences and utilizes local behavior similarity (LBS) to better represent sample relationships. The features used are pseudo-position specific scoring matrix (PsePSSM) [43], global encoding (GE) [44], normalized Moreau–Broto autocorrelation (NMBAC) [45], and combined (concatenation of PsePSSM, GE, and NMBAC). A new edge weight calculation method considering label information and a local distribution parameter is introduced. Additionally, the authors employ multiple Laplacian regularizations to construct a multigraph model, making it less sensitive to neighborhood size.

#### LapSVM [46]

Authors in [46] propose a semi-supervised learning approach, called the Laplacian support vector machine (LapSVM), which can be used to perform classification. The LapSVM works on the traditional support vector machine (SVM) on which, the manifold regularization (contains the geometric information of labeled and unlabeled samples) is applied. We use the same features as used in MLapSVM [42], i.e. pseudo-position specific scoring matrix (PsePSSM) [43], global encoding (GE) [44], normalized Moreau–Broto autocorrelation (NMBAC) [45], and combined (concatination of PsePSSM, GE, and NMBAC), to generate the embeddings, which as used as input to the LapSVM approach (as done in [42]).

#### SeqVec [47]

Authors in [47] introduce a method that utilizes an ELMo (Embeddings from Language Models) [48] based architecture called SeqVec, which involves several levels of processing. It starts by padding input sequences with special tokens and then uses character convolutions to map amino acids to a fixed-length latent space. A bidirectional Long Short Term Memory (LSTM) layer processes the sequence sequentially, introducing context-specific information. Another LSTM layer predicts the next word based on previous words. The forward and backward passes are optimized independently during training. The output from the SeqVec architecture is the vector representation, which is then used as input to classical machine learning classifiers, such as SVM, Naive Bayes (NB), Multi-Layer Perceptron (MLP), K-Nearest Neighbors (KNN), Random Forest (RF), Logistic Regression (LR), and Decision Tree (DT), for binding prediction (binary classification).

#### PDBP-Fusion [49]

Authors in [49] propose a methodology called PDBP-Fusion for predicting DNA-binding proteins based solely on primary sequence data. This method combines Convolutional Neural Networks (CNN) to capture local features and Bidirectional Long Short-Term Memory Networks (Bi-LSTM) to capture long-term dependencies in the DNA sequences. The framework consists of several layers for Sequence Encoding, Local Feature Learning, Long-Term Context Learning, and Synthetic Prediction. The Sequence Encoding layer prepares the DNA sequences for processing. In this method, two encoding methods are used including One-hot encoding, and Word embedding encoding, which represents discrete variables as continuous vectors. In Local Feature Learning, a CNN is employed to detect functional domains in protein sequences. This layer includes convolution, batch-normalization, ReLU activation, and max-pooling operations. It can use either One-hot encoding or Word embedding encoding. Long-Term Context Learning is used to capture long-term dependencies. It involves a Bi-LSTM layer. While CNN captures local characteristics, Bi-LSTM focuses on the broader context of gene sequences. Finally, in Synthetic Prediction, the previous layers’ outputs are concatenated into a vector and passed through a fully connected layer. The architecture uses the sigmoid activation function and cross-entropy loss function. The final output represents the prediction of DNA-binding proteins.

### 4.3 Data Visualization

A popular visualization technique, named t-SNE [50], is employed to visualize the feature vectors generated by the SeqVec [47] method. The t-SNE has been widely used for the visualization of biological data [51,52]. Note that we use SeqVec because that is the only method (among the discussed methods in this paper) that generates the embeddings and is not an end-to-end method for classification. The t-SNE plots are shown in Figure 1 for PDB2272, PDB1075, and PDB186 Datasets. The main idea for reporting the t-SNE plots is to observe if there is a clear class separation between the positive and negative class samples in the datasets. A clear class separation could mean that the problem is too simple and not worth exploring and vice versa. We can observe in Figure 1 that both positive and negative samples overlap, showing that there is no clear decision boundary that exists in the data initially.

**Fig. 1:**
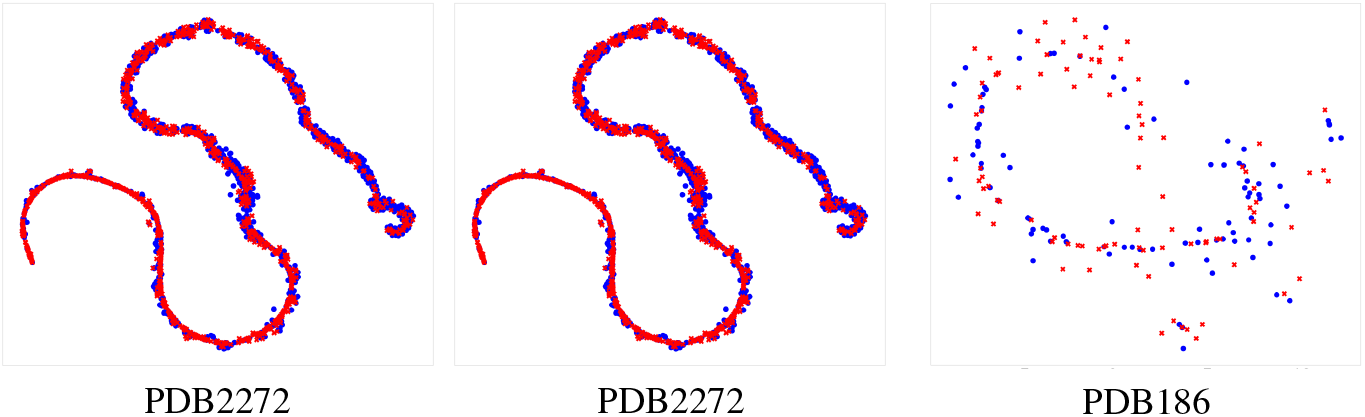
t-SNE Plots for SeqVec Embeddings for PDB2272, PDB1075, and PDB186 Datasets. The figure is best seen in color.

## 5 Results And Discussion

The results for the baselines and the proposed method are shown in Table 3 for the PDB14189 dataset on different evaluation metrics. Although the proposed method did not outperform the baselines, its performance is comparable for different evaluation metrics. One advantage that deep learning (DL) based architecture has, compared to the reported baselines, is the interpretability property. Since the simple ML classifiers, e.g. SVM, do not show great promise in terms of the explainability of results, the DL models are preferred, e.g. the proposed DeepPWM BindingNet, due to the inclusion of attention mechanism. Therefore, getting the highest predictive performance in this case is not the top priority, rather focusing on designing the architecture that shows comparable results while promising the architecture that holds promise for interpretability.

**Table 3:** Results comparison for PDB14189 dataset. The best values for each method are underlined.

| Method | Model | Acc. ↑ | Prec. ↑ | NPV ↑ | Sensitivity ↑ | Specificity ↑ | MCC ↑ | F1 ↑ | ROC-AUC ↑ | ROC-Pr ↑ |
| --- | --- | --- | --- | --- | --- | --- | --- | --- | --- | --- |
| Local Behavior Similarity (LapSVM) [46] | GE | 85.52 ± 0.20 | 83.83 ± 0.24 | 87.42 ± 0.75 | 88.19 ± 0.87 | 82.82 ± 0.47 | 71.13 ± 0.46 | 85.95 ± 0.29 | 92.85 ± 0.20 | 90.80 ± 0.19 |
| | NMBAC | 89.70 ± 0.01 | 85.29 ± 0.03 | 95.45 ± 0.03 | 96.07 ± 0.03 | 83.27 ± 0.04 | 80.04 ± 0.01 | 90.36 ± 0.01 | 95.93 ± 0.05 | 94.90 ± 0.04 |
| | MCD | 74.84 ± 0.29 | 73.48 ± 0.63 | 76.49 ± 1.49 | 78.16 ± 2.34 | 71.49 ± 1.77 | 49.81 ± 0.71 | 75.72 ± 0.76 | 82.45 ± 0.13 | 80.67 ± 0.02 |
| | PSSM | 76.88 ± 0.83 | 71.91 ± 1.52 | 85.19 ± 1.13 | 88.76 ± 1.56 | 64.89 ± 3.24 | 55.34 ± 1.06 | 79.42 ± 0.30 | 86.42 ± 0.09 | 84.50 ± 0.50 |
| | Combined | 74.00 ± 0.08 | 71.38 ± 0.05 | 77.43 ± 0.12 | 80.54 ± 0.13 | 67.39 ± 0.03 | 48.37 ± 0.17 | 75.69 ± 0.09 | 82.11 ± 0.17 | 81.98 ± 0.07 |
| Local Behavior Similarity (MLapSVM) [42] | GE | 74.64 ± 0.40 | 72.57 ± 0.76 | 77.19 ± 0.28 | 79.63 ± 0.66 | 69.59 ± 1.38 | 49.49 ± 0.73 | 75.93 ± 0.20 | 82.39 ± 0.40 | 82.26 ± 0.66 |
| | NMBAC | 74.07 ± 0.71 | 67.12 ± 0.61 | 91.14 ± 1.36 | 94.88 ± 0.88 | 53.06 ± 1.31 | 52.85 ± 1.47 | 78.62 ± 0.55 | 87.08 ± 0.40 | 85.20 ± 0.42 |
| | MCD | 76.99 ± 0.83 | 77.22 ± 0.36 | 76.80 ± 1.48 | 76.88 ± 2.13 | 77.10 ± 0.73 | 54.00 ± 1.62 | 77.04 ± 1.14 | 84.40 ± 0.75 | 82.72 ± 0.94 |
| | PSSM | 91.27 ± 0.33 | 88.11 ± 0.65 | 95.07 ± 0.58 | 95.53 ± 0.58 | 86.97 ± 0.85 | 82.83 ± 0.62 | 91.66 ± 0.29 | 96.50 ± 0.40 | 95.57 ± 0.72 |
| | Combined | 87.39 ± 0.50 | 85.71 ± 0.84 | 89.29 ± 0.70 | 89.91 ± 0.79 | 84.84 ± 1.08 | 74.88 ± 0.99 | 87.75 ± 0.46 | 93.93 ± 0.53 | 92.05 ± 1.06 |
| SeqVec [47] | SVM | 71.45 ± 0.01 | 71.51 ± 0.01 | 71.74 ± 0.01 | 72.31 ± 0.01 | 69.12 ± 0.02 | 42.42 ± 0.01 | 71.17 ± 0.01 | 71.81 ± 0.01 | 67.62 ± 0.01 |
| | NB | 55.11 ± 0.01 | 65.41 ± 0.02 | 78.45 ± 0.03 | 96.21 ± 0.01 | 13.90 ± 0.01 | 17.87 ± 0.02 | 45.14 ± 0.01 | 55.46 ± 0.00 | 52.85 ± 0.00 |
| | MLP | 74.43 ± 0.01 | 74.56 ± 0.01 | 74.38 ± 0.02 | 74.12 ± 0.03 | 75.66 ± 0.01 | 48.97 ± 0.01 | 74.32 ± 0.01 | 74.66 ± 0.01 | 69.32 ± 0.01 |
| | KNN | 72.57 ± 0.00 | 72.62 ± 0.00 | 72.88 ± 0.01 | 71.92 ± 0.01 | 73.77 ± 0.01 | 44.16 ± 0.01 | 72.73 ± 0.00 | 72.54 ± 0.00 | 69.51 ± 0.01 |
| | RF | 76.75 ± 0.00 | 76.74 ± 0.00 | 77.94 ± 0.00 | 78.17 ± 0.01 | 73.65 ± 0.01 | 52.31 ± 0.01 | 76.65 ± 0.00 | 76.11 ± 0.00 | 68.32 ± 0.01 |
| | LR | 72.54 ± 0.00 | 72.76 ± 0.00 | 73.99 ± 0.01 | 74.14 ± 0.01 | 70.11 ± 0.01 | 44.84 ± 0.01 | 72.95 ± 0.00 | 72.81 ± 0.00 | 67.56 ± 0.01 |
| | DT | 65.54 ± 0.00 | 65.11 ± 0.00 | 65.84 ± 0.01 | 64.33 ± 0.01 | 66.74 ± 0.01 | 31.69 ± 0.00 | 65.57 ± 0.00 | 65.30 ± 0.00 | 67.56 ± 0.00 |
| PDBP-Fusion [49] | 2 Layer CNN (OH) | 80.34 ± 0.89 | 81.64 ± 1.89 | 79.46 ± 2.93 | 78.75 ± 4.55 | 81.95 ± 3.15 | 60.90 ± 1.64 | 80.04 ± 1.62 | 88.74 ± 0.50 | 86.69 ± 0.85 |
| | 3 Layer CNN (OH) | 81.64 ± 1.38 | 80.98 ± 3.38 | 83.41 ± 4.68 | 83.61 ± 7.01 | 79.66 ± 5.80 | 63.82 ± 2.28 | 81.95 ± 2.35 | 90.01 ± 0.63 | 88.57 ± 0.88 |
| | Fusion (Embed) | 53.35 ± 3.07 | 64.24 ± 8.87 | 41.52 ± 21.24 | 38.89 ± 37.31 | 67.95 ± 39.57 | 9.72 ± 7.38 | 36.27 ± 22.55 | 56.74 ± 7.05 | 56.20 ± 5.41 |
| | Fusion (OH) | 80.49 ± 1.61 | 77.66 ± 4.03 | 85.35 ± 4.15 | 86.79 ± 5.37 | 74.13 ± 7.67 | 61.94 ± 2.25 | 81.72 ± 1.10 | 89.25 ± 0.72 | 87.66 ± 0.98 |
| DeepPWM-BindingNet | - | 79.80 ± 0.94 | 77.53 ± 1.65 | 82.69 ± 1.75 | 84.32 ± 2.29 | 75.24 ± 2.71 | 59.89 ± 1.87 | 80.74 ± 0.90 | 87.69 ± 0.88 | 86.44 ± 0.94 |

The results for the PDB2272 dataset are shown in Table 4. We can observe that in terms of average accuracy, the proposed method shows value in the top 5% accuracy. For the remaining evaluation metrics, although the performance of the proposed method is not the highest, it shows comparable results.

**Table 4:**
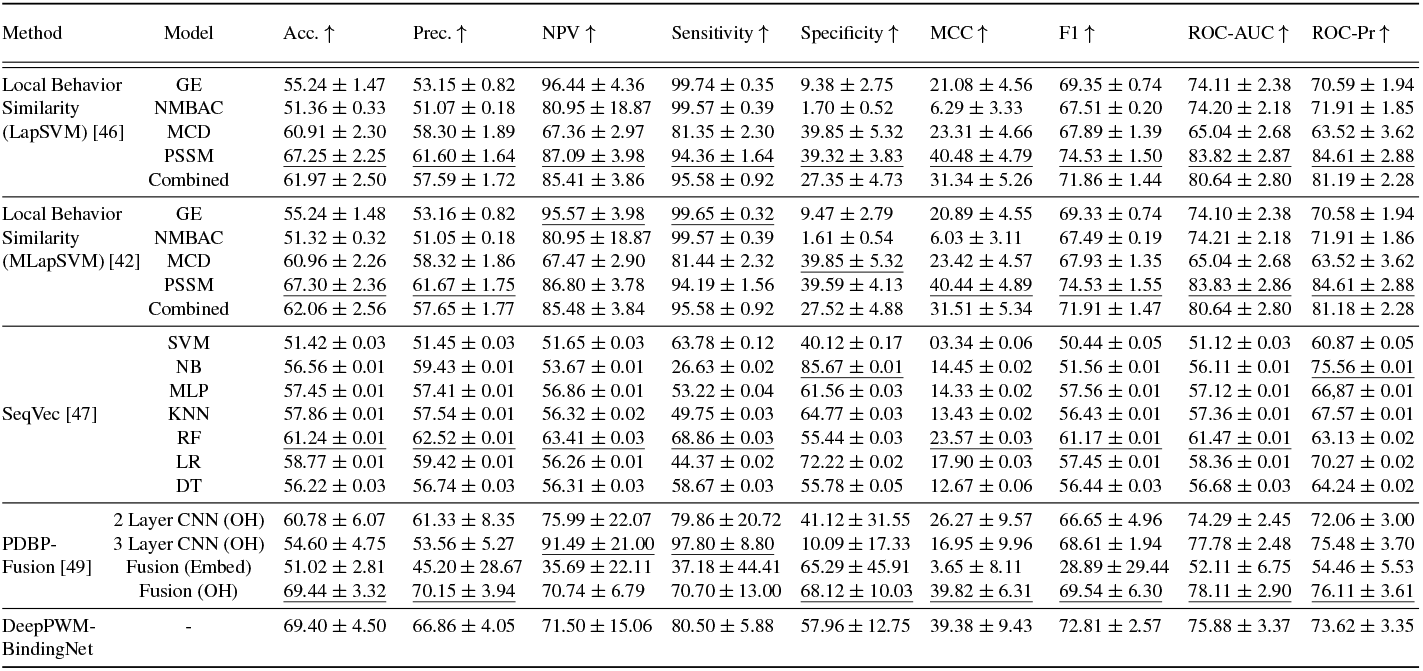
Results comparison for PDB2272 dataset. The best values for each method are underlined.

| Method | Model | Acc. ↑ | Prec. ↑ | NPV ↑ | Sensitivity ↑ | Specificity ↑ | MCC ↑ | F1 ↑ | ROC-AUC ↑ | ROC-Pr ↑ |
| --- | --- | --- | --- | --- | --- | --- | --- | --- | --- | --- |
| Local Behavior Similarity (LapSVM) [46] | GE | 55.24 ± 1.47 | 53.15 ± 0.82 | 96.44 ± 4.36 | 99.74 ± 0.35 | 9.38 ± 2.75 | 21.08 ± 4.56 | 69.35 ± 0.74 | 74.11 ± 2.38 | 70.59 ± 1.94 |
| | NMBAC | 51.36 ± 0.33 | 51.07 ± 0.18 | 80.95 ± 18.87 | 99.57 ± 0.39 | 1.70 ± 0.52 | 6.29 ± 3.33 | 67.51 ± 0.20 | 74.20 ± 2.18 | 71.91 ± 1.85 |
| | MCD | 60.91 ± 2.30 | 58.30 ± 1.89 | 67.36 ± 2.97 | 81.35 ± 2.30 | 39.85 ± 5.32 | 23.31 ± 4.66 | 67.89 ± 1.39 | 65.04 ± 2.68 | 63.52 ± 3.62 |
| | PSSM | 67.25 ± 2.25 | 61.60 ± 1.64 | 87.09 ± 3.98 | 94.36 ± 1.64 | 39.32 ± 3.83 | 40.48 ± 4.79 | 74.53 ± 1.50 | 83.82 ± 2.87 | 84.61 ± 2.88 |
| | Combined | 61.97 ± 2.50 | 57.59 ± 1.72 | 85.41 ± 3.86 | 95.58 ± 0.92 | 27.35 ± 4.73 | 31.34 ± 5.26 | 71.86 ± 1.44 | 80.64 ± 2.80 | 81.19 ± 2.28 |
| Local Behavior Similarity (MLapSVM) [42] | GE | 55.24 ± 1.48 | 53.16 ± 0.82 | 95.57 ± 3.98 | 99.65 ± 0.32 | 9.47 ± 2.79 | 20.89 ± 4.55 | 69.33 ± 0.74 | 74.10 ± 2.38 | 70.58 ± 1.94 |
| | NMBAC | 51.32 ± 0.32 | 51.05 ± 0.18 | 80.95 ± 18.87 | 99.57 ± 0.39 | 1.61 ± 0.54 | 6.03 ± 3.11 | 67.49 ± 0.19 | 74.21 ± 2.18 | 71.91 ± 1.86 |
| | MCD | 60.96 ± 2.26 | 58.32 ± 1.86 | 67.47 ± 2.90 | 81.44 ± 2.32 | 39.85 ± 5.32 | 23.42 ± 4.57 | 67.93 ± 1.35 | 65.04 ± 2.68 | 63.52 ± 3.62 |
| | PSSM | 67.30 ± 2.36 | 61.67 ± 1.75 | 86.80 ± 3.78 | 94.19 ± 1.56 | 39.59 ± 4.13 | 40.44 ± 4.89 | 74.53 ± 1.55 | 83.83 ± 2.86 | 84.61 ± 2.88 |
| | Combined | 62.06 ± 2.56 | 57.65 ± 1.77 | 85.48 ± 3.84 | 95.58 ± 0.92 | 27.52 ± 4.88 | 31.51 ± 5.34 | 71.91 ± 1.47 | 80.64 ± 2.80 | 81.18 ± 2.28 |
| SeqVec [47] | SVM | 51.42 ± 0.03 | 51.45 ± 0.03 | 51.65 ± 0.03 | 63.78 ± 0.12 | 40.12 ± 0.17 | 03.34 ± 0.06 | 50.44 ± 0.05 | 51.12 ± 0.03 | 60.87 ± 0.05 |
| | NB | 56.56 ± 0.01 | 59.43 ± 0.01 | 53.67 ± 0.01 | 26.63 ± 0.02 | 85.67 ± 0.01 | 14.45 ± 0.02 | 51.56 ± 0.01 | 56.11 ± 0.01 | 75.56 ± 0.01 |
| | MLP | 57.45 ± 0.01 | 57.41 ± 0.01 | 56.86 ± 0.01 | 53.22 ± 0.04 | 61.56 ± 0.03 | 14.33 ± 0.02 | 57.56 ± 0.01 | 57.12 ± 0.01 | 66.87 ± 0.01 |
| | KNN | 57.86 ± 0.01 | 57.54 ± 0.01 | 56.32 ± 0.02 | 49.75 ± 0.03 | 64.77 ± 0.03 | 13.43 ± 0.02 | 56.43 ± 0.01 | 57.36 ± 0.01 | 67.57 ± 0.01 |
| | RF | 61.24 ± 0.01 | 62.52 ± 0.01 | 63.41 ± 0.03 | 68.86 ± 0.03 | 55.44 ± 0.03 | 23.57 ± 0.03 | 61.17 ± 0.01 | 61.47 ± 0.01 | 63.13 ± 0.02 |
| | LR | 58.77 ± 0.01 | 59.42 ± 0.01 | 56.26 ± 0.01 | 44.37 ± 0.02 | 72.22 ± 0.02 | 17.90 ± 0.03 | 57.45 ± 0.01 | 58.36 ± 0.01 | 70.27 ± 0.02 |
| | DT | 56.22 ± 0.03 | 56.74 ± 0.03 | 56.31 ± 0.03 | 58.67 ± 0.03 | 55.78 ± 0.05 | 12.67 ± 0.06 | 56.44 ± 0.03 | 56.68 ± 0.03 | 64.24 ± 0.02 |
| PDBP-Fusion [49] | 2 Layer CNN (OH) | 60.78 ± 6.07 | 61.33 ± 8.35 | 75.99 ± 22.07 | 79.86 ± 20.72 | 41.12 ± 31.55 | 26.27 ± 9.57 | 66.65 ± 4.96 | 74.29 ± 2.45 | 72.06 ± 3.00 |
| | 3 Layer CNN (OH) | 54.60 ± 4.75 | 53.56 ± 5.27 | 91.49 ± 21.00 | 97.80 ± 8.80 | 10.09 ± 17.33 | 16.95 ± 9.96 | 68.61 ± 1.94 | 77.78 ± 2.48 | 75.48 ± 3.70 |
| | Fusion (Embed) | 51.02 ± 2.81 | 45.20 ± 28.67 | 35.69 ± 22.11 | 37.18 ± 44.41 | 65.29 ± 45.91 | 3.65 ± 8.11 | 28.89 ± 29.44 | 52.11 ± 6.75 | 54.46 ± 5.53 |
| | Fusion (OH) | 69.44 ± 3.32 | 70.15 ± 3.94 | 70.74 ± 6.79 | 70.70 ± 13.00 | 68.12 ± 10.03 | 39.82 ± 6.31 | 69.54 ± 6.30 | 78.11 ± 2.90 | 76.11 ± 3.61 |
| DeepPWM-BindingNet | - | 69.40 ± 4.50 | 66.86 ± 4.05 | 71.50 ± 15.06 | 80.50 ± 5.88 | 57.96 ± 12.75 | 39.38 ± 9.43 | 72.81 ± 2.57 | 75.88 ± 3.37 | 73.62 ± 3.35 |

The results for the PDB1075 dataset are shown in Table 5 for the proposed and baseline methods. We can observe that the proposed method is among the top 5% in terms of binding prediction using the majority of the evaluation metrics.

**Table 5:** Results comparison for PDB1075 dataset. The best values for each method are underlined.

| Method | Model | Acc. ↑ | Prec. ↑ | NPV ↑ | Sensitivity ↑ | Specificity ↑ | MCC ↑ | F1 ↑ | ROC-AUC ↑ | ROC-Pr ↑ |
| --- | --- | --- | --- | --- | --- | --- | --- | --- | --- | --- |
| Local Behavior Similarity (LapSVM) [46] | GE | 51.16 ± 0.00 | 0.00 ± 0.00 | 51.16 ± 0.00 | 0.00 ± 0.00 | <u>100.00 ± 0.00</u> | 0.00 ± 0.00 | 0.00 ± 0.00 | 77.50 ± 2.46 | 74.97 ± 1.55 |
| | NMBAC | 51.16 ± 0.00 | 0.00 ± 0.00 | 51.16 ± 0.00 | 0.00 ± 0.00 | <u>100.00 ± 0.00</u> | 0.00 ± 0.00 | 0.00 ± 0.00 | 77.11 ± 4.24 | 74.41 ± 4.08 |
| | MCD | 61.21 ± 1.37 | <u>81.29 ± 5.55</u> | 57.41 ± 0.89 | 27.05 ± 3.16 | 93.82 ± 2.47 | 28.35 ± 3.58 | 40.41 ± 3.51 | 76.20 ± 1.78 | 73.95 ± 2.87 |
| | PSSM | <u>75.07 ± 4.55</u> | 80.06 ± 5.50 | 71.84 ± 4.08 | <u>65.14 ± 5.83</u> | 84.55 ± 4.11 | <u>50.78 ± 9.23</u> | <u>71.79 ± 5.49</u> | <u>83.06 ± 3.58</u> | <u>79.57 ± 4.49</u> |
| | Combined | 70.70 ± 3.87 | 78.79 ± 5.30 | 66.70 ± 3.43 | 54.86 ± 6.64 | 85.82 ± 4.13 | 43.00 ± 7.84 | 64.48 ± 5.62 | 80.55 ± 3.34 | 76.41 ± 4.46 |
| Local Behavior Similarity (MLapSVM) [42] | GE | 51.16 ± 0.00 | 0.00 ± 0.00 | 51.16 ± 0.00 | 0.00 ± 0.00 | <u>100.00 ± 0.00</u> | 0.00 ± 0.00 | 0.00 ± 0.00 | 77.49 ± 2.45 | 74.98 ± 1.53 |
| | NMBAC | 51.07 ± 0.19 | 0.00 ± 0.00 | 51.12 ± 0.09 | 0.00 ± 0.00 | 99.82 ± 0.36 | -1.34 ± 2.67 | 0.00 ± 0.00 | 77.11 ± 4.24 | 74.41 ± 4.08 |
| | MCD | 61.12 ± 1.23 | 81.21 ± 5.48 | 57.34 ± 0.80 | 26.86 ± 3.04 | 93.82 ± 2.47 | 28.17 ± 3.34 | 40.18 ± 3.30 | 76.21 ± 1.77 | 73.94 ± 2.87 |
| | PSSM | 74.88 ± 4.44 | 79.98 ± 5.45 | 71.61 ± 3.97 | 64.76 ± 5.68 | 84.55 ± 4.11 | 50.44 ± 9.02 | 71.52 ± 5.35 | 83.07 ± 3.60 | 79.58 ± 4.50 |
| | Combined | 70.42 ± 3.49 | 78.98 ± 5.18 | 66.29 ± 2.99 | 53.90 ± 5.83 | 86.18 ± 4.04 | 42.58 ± 7.17 | 63.90 ± 5.00 | 80.56 ± 3.33 | 76.42 ± 4.45 |
| SeqVec [47] | SVM | 48.56 ± 0.05 | 48.43 ± 0.05 | 50.26 ± 0.05 | 37.23 ± 0.11 | 59.98 ± 0.05 | -4.11 ± 0.10 | 47.12 ± 0.05 | 48.45 ± 0.05 | 63.32 ± 0.02 |
| | NB | 57.45 ± 0.04 | 60.67 ± 0.04 | 66.43 ± 0.04 | 80.22 ± 0.03 | 36.56 ± 0.06 | 18.88 ± 0.07 | 55.26 ± 0.05 | 58.68 ± 0.03 | 56.89 ± 0.03 |
| | MLP | 52.87 ± 0.02 | 52.89 ± 0.03 | 54.55 ± 0.04 | 55.93 ± 0.05 | 50.72 ± 0.05 | 4.45 ± 0.05 | 52.57 ± 0.02 | 52.32 ± 0.03 | 61.45 ± 0.02 |
| | KNN | 58.56 ± 0.02 | 58.67 ± 0.03 | 60.66 ± 0.03 | 62.35 ± 0.06 | 54.88 ± 0.03 | 16.43 ± 0.05 | 58.32 ± 0.02 | 58.56 ± 0.02 | 62.22 ± 0.02 |
| | RF | 61.78 ± 0.02 | 61.43 ± 0.02 | 61.22 ± 0.03 | 55.56 ± 0.04 | 67.67 ± 0.03 | 22.78 ± 0.04 | 61.32 ± 0.02 | 61.45 ± 0.02 | 66.57 ± 0.02 |
| | LR | 53.32 ± 0.04 | 53.45 ± 0.04 | 56.38 ± 0.05 | 62.92 ± 0.06 | 44.58 ± 0.02 | 6.59 ± 0.08 | 52.26 ± 0.03 | 53.43 ± 0.04 | 59.44 ± 0.01 |
| | DT | 57.67 ± 0.03 | 57.55 ± 0.03 | 59.43 ± 0.02 | 57.81 ± 0.04 | 57.99 ± 0.06 | 15.64 ± 0.06 | 57.32 ± 0.03 | 57.67 ± 0.03 | 63.28 ± 0.03 |
| PDBP-Fusion [49] | 2 Layer CNN (OH) | <u>68.65 ± 2.86</u> | <u>66.72 ± 5.99</u> | 75.58 ± 8.05 | 75.73 ± 15.14 | 61.96 ± 15.27 | 39.85 ± 5.06 | 69.45 ± 5.47 | 78.08 ± 2.41 | 74.29 ± 2.97 |
| | 3 Layer CNN (OH) | 68.33 ± 3.69 | 62.96 ± 4.57 | 82.47 ± 5.29 | 87.64 ± 7.20 | 50.15 ± 12.33 | 41.27 ± 5.33 | 72.87 ± 1.94 | 79.25 ± 1.78 | 76.38 ± 2.72 |
| | Fusion (Embed) | 51.10 ± 3.27 | 28.06 ± 28.71 | 40.51 ± 23.96 | 45.46 ± 47.69 | 56.40 ± 46.73 | 0.63 ± 8.38 | 31.31 ± 32.45 | 68.03 ± 5.07 | 66.48 ± 5.64 |
| | Fusion (OH) | 65.04 ± 5.67 | 59.57 ± 4.87 | <u>86.05 ± 6.42</u> | <u>92.01 ± 6.47</u> | 39.64 ± 14.98 | 37.15 ± 9.32 | 71.96 ± 2.82 | <u>80.09 ± 1.67</u> | <u>76.92 ± 3.31</u> |
| DeepPWM-BindingNet | - | 72.05 ± 2.56 | 69.37 ± 3.48 | 75.66 ± 3.54 | 76.37 ± 5.43 | 68.00 ± 6.19 | 44.70 ± 5.03 | 72.52 ± 2.57 | 78.45 ± 2.70 | 74.13 ± 5.23 |

The binding prediction results for the PDB186 dataset are reported in Table 6 for the proposed and baseline models. Although the SVM-based baselines show higher predictive performance, the proposed method shows a near-perfect score for the sensitivity metric. Moreover, it shows a comparable performance for F1 and ROC compared to the baselines.

**Table 6:** Results comparison for PDB186 dataset. The best values for each method are underlined.

| Method | Model | Acc. ↑ | Prec. ↑ | NPV ↑ | Sensitivity ↑ | Specificity ↑ | MCC ↑ | F1 ↑ | ROC-AUC ↑ | ROC-Pr ↑ |
| --- | --- | --- | --- | --- | --- | --- | --- | --- | --- | --- |
| Local Behavior Similarity (LapSVM) [46] | GE | 52.70 ± 5.92 | 40.71 ± 21.36 | 56.03 ± 8.71 | 54.80 ± 36.51 | 50.99 ± 30.86 | 6.36 ± 12.92 | 45.22 ± 26.86 | 61.03 ± 8.05 | 59.99 ± 6.81 |
| | NMBAC | 49.47 ± 1.60 | 0.00 ± 0.00 | 49.73 ± 1.32 | 0.00 ± 0.00 | <u>98.95 ± 2.11</u> | -3.29 ± 6.58 | 0.00 ± 0.00 | 68.62 ± 6.32 | 65.23 ± 6.79 |
| | MCD | 53.23 ± 3.15 | 55.25 ± 7.51 | 52.24 ± 1.82 | 34.15 ± 10.88 | 71.99 ± 11.22 | 6.77 ± 8.10 | 41.21 ± 9.53 | 57.49 ± 4.36 | 60.72 ± 5.85 |
| | PSSM | <u>66.15 ± 7.08</u> | <u>66.08 ± 9.23</u> | 67.61 ± 6.23 | 69.71 ± 9.35 | 62.40 ± 14.50 | <u>32.87 ± 14.11</u> | 67.31 ± 6.61 | 70.15 ± 8.44 | 71.77 ± 6.48 |
| | Combined | 65.59 ± 11.00 | 63.71 ± 11.10 | <u>69.53 ± 12.18</u> | <u>73.98 ± 13.07</u> | 57.08 ± 14.29 | 32.09 ± 22.01 | 68.16 ± 10.85 | 67.32 ± 8.77 | 67.99 ± 5.46 |
| Local Behavior Similarity (MLapSVM) [42] | GE | 52.70 ± 5.92 | 40.71 ± 21.36 | 56.03 ± 8.71 | 54.80 ± 36.51 | 50.99 ± 30.86 | 6.36 ± 12.92 | 45.22 ± 26.86 | 61.09 ± 8.03 | 60.10 ± 6.72 |
| | NMBAC | 49.47 ± 1.60 | 0.00 ± 0.00 | 49.73 ± 1.32 | 0.00 ± 0.00 | <u>98.95 ± 2.11</u> | -3.29 ± 6.58 | 0.00 ± 0.00 | 68.68 ± 6.29 | 65.26 ± 6.78 |
| | MCD | 53.23 ± 3.15 | 55.25 ± 7.51 | 52.24 ± 1.82 | 34.15 ± 10.88 | 71.99 ± 11.22 | 6.77 ± 8.10 | 41.21 ± 9.53 | 57.43 ± 4.33 | 60.60 ± 5.74 |
| | PSSM | <u>65.60 ± 6.40</u> | <u>65.80 ± 8.81</u> | 66.98 ± 5.83 | 68.60 ± 9.96 | 62.40 ± 14.50 | 31.85 ± 12.83 | 66.51 ± 6.13 | <u>70.10 ± 8.36</u> | <u>71.72 ± 6.41</u> |
| | Combined | 65.59 ± 11.00 | 63.71 ± 11.10 | <u>69.53 ± 12.18</u> | <u>73.98 ± 13.07</u> | 57.08 ± 14.29 | 32.09 ± 22.01 | 68.16 ± 10.85 | 67.31 ± 8.78 | 67.98 ± 5.44 |
| SeqVec [47] | SVM | 50.11 ± 0.04 | 49.25 ± 0.05 | 48.32 ± 0.11 | 53.91 ± 0.12 | 46.45 ± 0.20 | -1.54 ± 0.11 | 49.57 ± 0.05 | 50.43 ± 0.05 | 61.56 ± 0.04 |
| | NB | 60.43 ± 0.05 | 61.61 ± 0.04 | 62.67 ± 0.04 | 65.17 ± 0.08 | 56.57 ± 0.11 | 21.52 ± 0.09 | 60.12 ± 0.05 | 60.76 ± 0.05 | 63.43 ± 0.04 |
| | MLP | 50.32 ± 0.06 | 51.64 ± 0.06 | 52.87 ± 0.09 | 54.43 ± 0.16 | 48.32 ± 0.13 | 2.43 ± 0.12 | 50.41 ± 0.06 | 51.89 ± 0.06 | 61.65 ± 0.05 |
| | KNN | 56.24 ± 0.06 | 58.46 ± 0.06 | 57.67 ± 0.09 | 52.55 ± 0.15 | <u>63.57 ± 0.17</u> | 15.13 ± 0.12 | 56.92 ± 0.07 | 57.59 ± 0.06 | <u>66.26 ± 0.08</u> |
| | RF | 53.35 ± 0.04 | 54.15 ± 0.03 | 54.23 ± 0.06 | 51.15 ± 0.13 | 55.73 ± 0.12 | 7.91 ± 0.07 | 52.35 ± 0.04 | 53.91 ± 0.03 | 63.55 ± 0.05 |
| | LR | 51.46 ± 0.08 | 52.32 ± 0.08 | 53.57 ± 0.11 | 52.68 ± 0.12 | 51.43 ± 0.07 | 3.57 ± 0.16 | 51.79 ± 0.08 | 52.32 ± 0.08 | 62.46 ± 0.02 |
| | DT | 49.45 ± 0.02 | 50.43 ± 0.03 | 50.42 ± 0.04 | 48.56 ± 0.14 | 51.78 ± 0.14 | -1.48 ± 0.06 | 48.36 ± 0.02 | 50.78 ± 0.03 | 62.47 ± 0.06 |
| PDBP-Fusion [49] | 2 Layer CNN (OH) | 56.20 ± 6.36 | 55.71 ± 5.94 | 54.67 ± 30.92 | 79.31 ± 22.77 | 33.54 ± 27.62 | <u>15.38 ± 13.17</u> | 62.89 ± 10.17 | 65.30 ± 5.84 | 67.28 ± 5.55 |
| | 3 Layer CNN (OH) | 51.83 ± 4.26 | 51.20 ± 3.14 | 22.07 ± 30.17 | 95.18 ± 8.03 | 8.34 ± 14.15 | 4.20 ± 9.43 | 66.35 ± 2.67 | 64.12 ± 5.21 | 64.52 ± 5.35 |
| | Fusion (Embed) | 50.86 ± 1.83 | 40.33 ± 28.42 | 22.27 ± 28.85 | 64.43 ± 47.44 | <u>36.63 ± 47.62</u> | 2.50 ± 7.05 | 43.78 ± 31.24 | 65.37 ± 7.42 | 67.24 ± 5.53 |
| | Fusion (OH) | 53.55 ± 6.67 | 52.70 ± 5.40 | 24.46 ± 33.68 | 93.31 ± 10.97 | <u>13.81 ± 20.83</u> | 8.01 ± 14.64 | <u>66.75 ± 3.58</u> | <u>67.31 ± 5.57</u> | <u>69.13 ± 4.90</u> |
| DeepPWM-BindingNet | - | 49.89 ± 2.81 | 50.02 ± 2.22 | 6.55 ± 17.86 | 97.61 ± 5.05 | 2.20 ± 7.83 | -1.58 ± 7.77 | 66.05 ± 1.93 | 60.00 ± 11.82 | 64.34 ± 8.68 |

Overall, while our proposed approach may not have surpassed the baselines in terms of raw performance metrics in some cases, its unique characteristics and advantages in addressing specific challenges within the problem domain make it a compelling addition to the field. By venturing into previously uncharted territory, our method has opened up new avenues of exploration and has the potential to provide robust and versatile solutions. Moreover, its resource efficiency, interpretability, and adaptability offer practical benefits that cannot be overlooked. We acknowledge that in complex domains, no single method may excel in all aspects, but our approach, by complementing existing techniques (e.g. the idea of PWM) and offering smoother learning from the data, contributes to a more comprehensive toolkit for researchers and practitioners. Furthermore, its potential for improved real-world applicability and ethical considerations position it as a promising foundation for future research endeavors. While it may not be the ultimate panacea, our proposed approach brings forth advantages and insights that enrich the field and prompt exciting directions for further exploration. Also, showing results using different types of evaluation metrics for the popular benchmark datasets (which, to the best of our knowledge, is not reported to this extent in the literature) provides a comprehensive analysis of the proposed and baseline methods, which researchers can use as a benchmark for extended studies.

### 5.1 Limitations and Future Work

The limitation of the proposed approach is its computational cost as it demands higher resources, and takes longer to compute features. Another limitation of this work is using a single type of data (i.e. DNA-binding prediction). Future work will explore other novel ideas in deep learning, such as transfer learning. Moreover, applying the proposed method to other biological applications could show the generalizability of the model.

## 6 Conclusion

In this work, we introduced DeepPWM-BindingNet, a novel deep-learning framework tailored for the prediction of DNA-protein binding interactions. This framework effectively addresses the crucial challenge of accurately identifying binding sites, with significant implications for molecular biology, gene regulation, and genome functionality research. Our approach leverages the rich information encoded in Position Weight Matrices, which capture the sequence-specific binding preferences of proteins. By seamlessly integrating PWM-derived features into a hybrid model of convolutional recurrent neural networks (CRNNs), we achieve a comprehensive representation of DNA sequences and protein structures. This hierarchical feature extraction process enables the model to capture both local and global sequence patterns, enhancing its predictive accuracy. Moreover, we introduced an attention mechanism that allows the model to focus on critical regions within sequences. This mechanism improves the model’s capacity to locate essential binding sites and also enhances its interpretability. Researchers can gain deeper insights into the underlying binding mechanisms, which is essential for advancing our understanding of gene regulation. The integration of PWM-derived features further boosts the model’s predictive ability, making it a valuable tool for deciphering intricate DNA-protein interactions. It offers a powerful and interpretable solution for predicting DNA-protein binding interactions, contributing to our comprehension of gene regulation mechanisms and opening new avenues for biological research.

## 7 Acknowldgement

Sarwan Ali, Prakash Chourasia, and Murray Patterson would like to thank Molecular Basis of Disease (MBD) and Georgia State University for their support and resources.

## Footnotes

1 http://www.uniprot.org/

2 http://www.ebi.ac.uk/swissprot/

